# Beyond Homology Transfer: Deep Learning for Automated Annotation of Proteins

**DOI:** 10.1101/168120

**Authors:** Mohammad Nauman, Hafeez Ur Rehman, Gianfranco Politano, Alfredo Benso

## Abstract

Accurate annotation of protein functions is important for a profound understanding of molecular biology. A large number of proteins remain uncharacterized because of the sparsity of available supporting information. For a large set of uncharacterized proteins, the only type of information available is their amino acid sequence. In this paper, we propose DeepSeq – a deep learning architecture – that utilizes only the protein sequence information to predict its associated functions. The prediction process does not require handcrafted features; rather, the architecture automatically extracts representations from the input sequence data. Results of our experiments with DeepSeq indicate significant improvements in terms of prediction accuracy when compared with other sequence-based methods. Our deep learning model achieves an overall validation accuracy of 86.72%, with an F1 score of 71.13%. Moreover, using the automatically learned features and without any changes to DeepSeq, we successfully solved a different problem i.e. protein function localization, with no human intervention. Finally, we discuss how this same architecture can be used to solve even more complicated problems such as prediction of 2D and 3D structure as well as protein-protein interactions.

## Introduction

A biological cell is an intricately arranged chemical factory, the sophisticated organization of which results into multi-cellular species with staggering complexities. At molecular level proteins are the main workhorses of these living cells. The knowledge of protein functions is of paramount importance for an ample understanding of these complex molecular machines that ultimately govern life. Accurate identification of protein function has implications in a wide variety of areas, which includes, discovering new therapeutic interventions, understanding novel diseases as well as designing their cures, better agriculture etc. Experimental methods are generally used to study protein functions but they cannot scale up to the task, due to associated cost and time. On the other hand, high throughput experiments like next generation sequencing technologies are resulting in a large number of new protein sequences that remain uncharacterized^1^. Figure 1 shows the latest major database statistics, which reflect a rapidly increasing gap between known sequences (GenBank and EMBL curves) and structurally characterized sequences (PDB curve). This tremendous growth of uncharacterized proteins poses a serious challenge at the forefront of molecular biology. To overcome this, computational approaches are generally relied upon for the annotation of protein functions.

**Figure 1.**
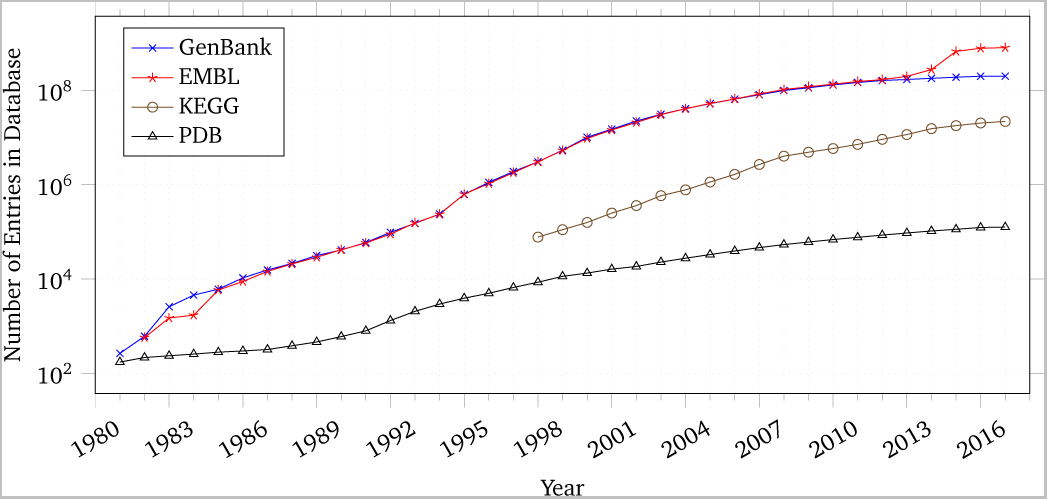
Growth of major sequence, pathway and 3D structure databases^1^. The amount of available sequence information (GenBank, EMBL) is orders of magnitude greater than that associated with structure information (PDB)

Many techniques have been proposed in the recent past that exploit a wide variety of data to predict protein functions^2^–^7^. A fundamental way to annotate proteins is through homology based association. The most established algorithms achieve the task of computational functional inference through homology by finding Basic Local Alignment Search Tool (BLAST) hits^8^, among proteins that have experimentally derived functions.

Many techniques utilize information derived from high throughput experiments that include protein complexes^7^, gene expression profiles^9^, protein-protein interactions (PPIs)^10^, protein structure data^11^, phylogenetic profiles^12^, and micro-arrays data sets^13^ etc. The most well-known techniques are based on proteome-scale PPI networks that are obtained from several organisms. These techniques utilize protein-protein interactions data in a variety of ways. The algorithms differs from each other by the way they utilize either the global properties or the topological properties of a multi-species interactome e.g.,^14^–^17^. Moreover, these methods also incorporate quite varied underlying formulations and construct models by using well understood concepts from the fields of graph theory, graphical models, stochastic processes, probabilistic graphs as well as clustering concepts e.g,^15, 17^. Integration of many types of biological information from heterogeneous biological sources has also proven to significantly increase the overall accuracy of function prediction techniques^2, 18, 19^, because each type of information gives separate clues and contributes towards the annotation of protein function.

Unfortunately, the sequence function gap keeps increasing^1^, and an extraordinary large number of proteins remain uncharacterized because they have very limited supporting information (e.g., 3D structure, co-complex localization etc.) available that may have been helpful in their computational annotation. The only information available for such proteins is their amino acid sequence. Thus, annotation of such proteins can be performed by utilizing sequence information only, e.g., through homology (i.e., sequence homology based function transfer). On the other hand, sequence homology face two major challenges in terms of its implication for annotation transfer. First, not all uncharacterized proteins have experimentally verified homologs available, thus limiting its application. Secondly, higher sequence similarity does not always guarantee same function i.e., a small perturbation in sequence information may force drastic changes in function. In other words, two sequences having, say, 99% sequence similarity may exhibit different functions^20^. Due to these two challenges, annotating uncharacterized proteins with sparse biological information is still an open research challenge and there is a dire need of algorithms that accurately annotate uncharacterized proteins with sparse supporting information i.e given only protein sequences. A protein sequence is a highly compact form of complex biological information.

The issue of extracting information from highly complex and unstructured sources has been seen repeatedly in several domains. Deep learning^21^ has been shown to be extremely effective in addressing issues of this nature. The novelty of deep learning as opposed to traditional machine learning pipelines is depicted in Figure 2. In classical machine learning, inputs are taken by human experts and features are extracted through low-level algorithms. In vision, for instance, SIFT^22^ is a popular feature extraction algorithm. Afterwards, these features are clustered and finally a machine learning classifier (such as Support Vector Machines^23^) is used to assign weights to the features. In essence, the machine is only learning to assign weights to hand-crafted features. On the other hand, in the paradigm of deep learning, the inputs are provided in their raw form to the learning pipeline which not only performs classification but also extracts features as part of its end-to-end learning. This means that the expertise of humans is not required during the whole learning phase thus reducing the time, complexity and efforts of the whole process significantly.

**Figure 2.**
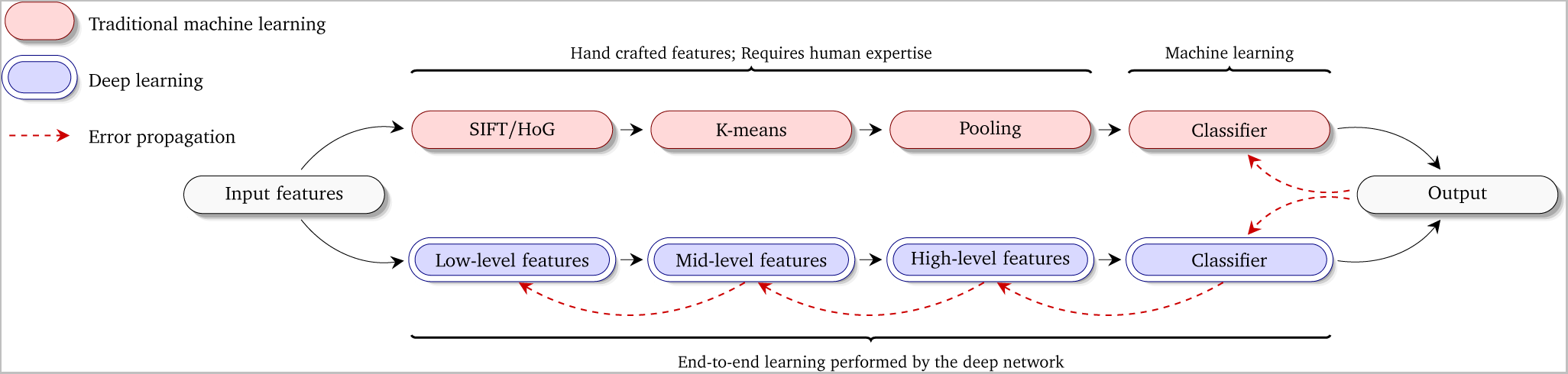
Traditional vs deep learning pipeline. In traditional machine learning, a lot of human time and effort goes into feature engineering. Deep learning, on the other hand, employs an end-to-end architecture that requires no human intervention for feature extraction.

Recent advances in deep learning provide an automated way of enabling machines to learn classification from raw data without the involvement of a human expert in the pipeline and has seen immense success in a myriad of fields including computer vision^24^, speech synthesis^25^, natural language processing^26^ and machine translation^27^ among several others. Results in these domains advocate for the fact that, through recently developed methods, machines can extract meaningful information from unstructured data far better than human experts, at a fraction of the cost and time.

In this paper, we propose and evaluate *DeepSeq* – a new deep learning architecture for protein function characterization. The fundamental conceptual innovation of our approach is to utilize the only available information about an uncharacterized protein i.e., its amino acid sequence. Importantly, DeepSeq does not require any hand crafted features for function prediction. We applied DeepSeq to the prediction of functions for Homo sapiens species proteins, using Gene Ontology (GO) as the annotation standard. The results indicate significant improvements in terms of prediction accuracy when compared with other sequence based methods.

In a nutshell, the contributions of our work are the following: (a) We design and develop a new deep learning architecture that is able to predict molecular functions given only the protein sequences as input (b) We use the same architecture to perform protein localization without human intervention or referring to homology information (c) We provide the code for the whole pipeline as open source at http://github.com/recluze/deepseq.^1^ The code is modular in nature so that our method can be verified and extended by varying the DeepSeq architecture and/or running experiments on sequences of other species.

## Materials and methods

In this section, we describe DeepSeq – the deep learning architecture we have developed to perform end-to-end learning of protein functions from sequences. We discuss the architecture that has achieved the best results in our studies along with the rationale for model parameters and hyperparameters.

### Model inputs

Our best model is primarily a convolutional neural network with tweaks that make it suitable for the particular problem of protein function prediction. A high-level view of our network architecture is shown in Figure 3. Inputs to the network are protein sequences with a maximum size of 2,000. We removed sequences larger than this threshold in order to reduce training times. However, predictions can be made using our resulting architecture for sequences of larger size as well. Smaller sequences were 0-padded before feeding them to the network. Hence, the size of inputs to our network is represented as (*ψ,* 2000), where *ψ* is the batch size. Smaller batch sizes lead to faster training at the cost of reduced accuracy. In our experiments, we set the batch size of 500.

**Figure 3.**
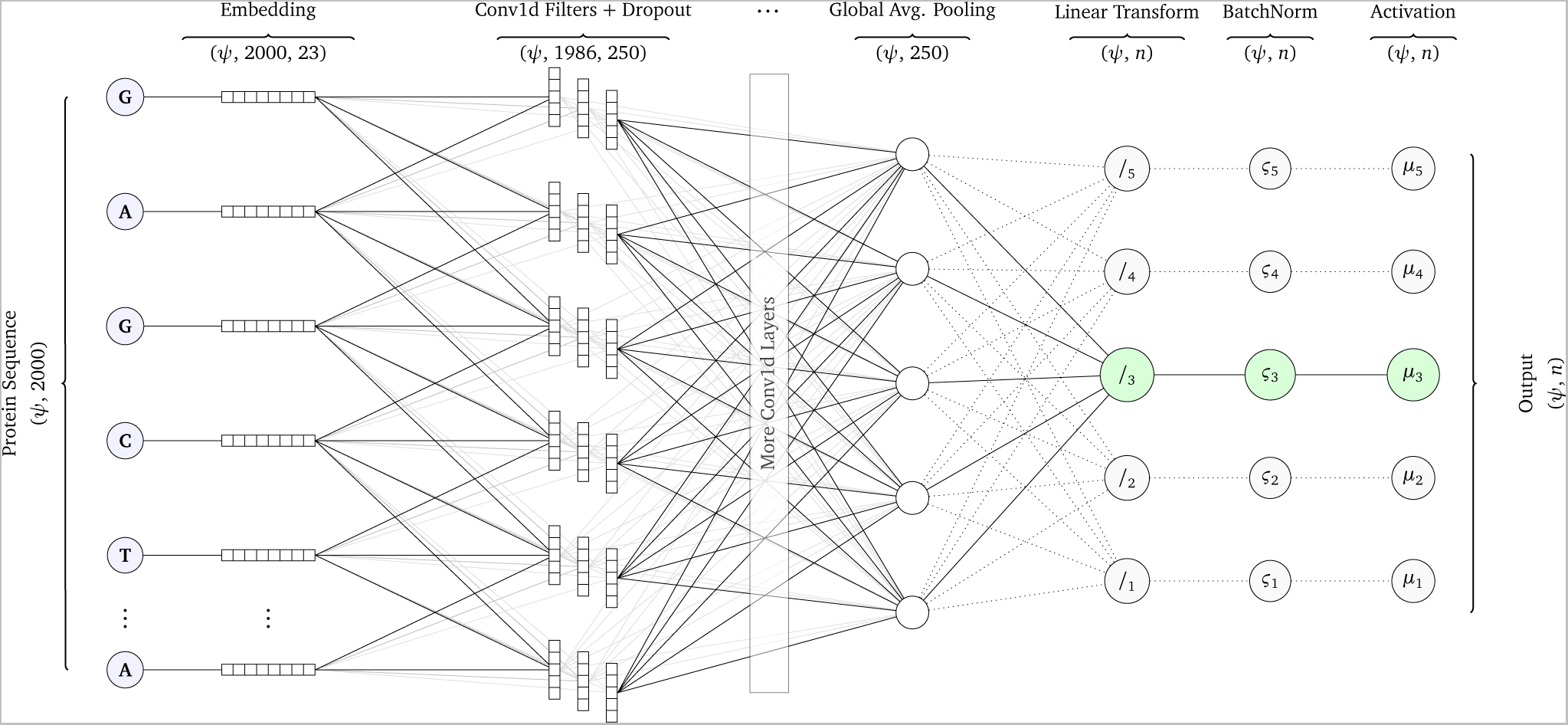
Network architecture for DeepSeq – our deep learning architecture for function prediction through protein sequences. DeepSeq is a true deep learning architecture since it employs end-to-end learning. Thus, the machine not only predicts protein functions but also learns to extract meaningful features automatically.

### Feature representation layers

Once the inputs have been fed to DeepSeq, it would extract features or representations from the inputs. Discrete and unordered inputs (i.e. amino acids in our case) are typically encoded as one-hot vectors^28^ and then fed to the neural network. However, one-hot representation embeds unique symbols equidistant from each other in a *l*-dimensional space where *l* is the number of unique values the symbol can take. In our case, there are 22 possible amino acids (plus a 0 for padding). A better alternative, that has shown to aid in learning, is to use a concept similar to word embeddings^29, 30^. Through embeddings, symbols that are similar to each other are represented with vectors that are closer in an *h*-dimensional space where *h ≤ l*. For our architecture, we set *h* equal to *l* thus each amino acid is represented as a 23-dimensional vector. Vectors closer to each other in this 23-dimensional space are similar. Notice that since the embedding layer is part of DeepSeq, this similarity is also learned automatically from data. This concept of learning all model parameters from data is the hallmark of end-to-end models and leads to much superior results^31^.

These embeddings are then passed to the layers responsible for transforming the data in a representation which can be discriminated. Since we have a fairly long sequence of amino acids, it is not possible to use a fully connected layer. The number of parameters required to train just a single layer would be close to 14 million for a small layer of just 200 nodes. This would lead to unfeasibly slow learning. We therefore chose to use sparse connections through convolutional layers. Our first sparse layer uses conv1d filters of length 15 and houses 250 filters while the second layer has 100 filters of the same length. Since weights are shared by the filters for different instantiations, this leads to a much smaller number of model parameters. To cater to the problem of overfitting that surfaced in our earlier experiments, we used a dropout layer with a dropout factor of 0.3 after each conv1d layer. Dropout^24^ is a highly popular form of regularization that not only improves the generalization of our algorithm but also reduces the time required to learn. The problem of vanishing gradients that slows down learning (or stops it altogether) was catered to by using the rectified linear unit activation^32^ in conv1d layers.

Typically, convolution layers are joined to the dense layers that perform classification through a flattening operation. However, again, due to the extremely large number of nodes in our model, flattening is simply not feasible. We therefore chose to use a global pooling operation which significantly reduces the number of parameters at the cost of slightly reduced accuracy. The final layer after this pooling operation is a single dense layer. For this complete architecture, the total number of trainable parameters is 462,234, which is significantly lesser than the naïve implementation of densely connected or even convolution layers. The output of these layers acts as the feature representations that DeepSeq has automatically extracted from the input sequence.

### Normalization and output layers

Based on the features extracted in the previous layers, a discriminator can be applied which performs the actual prediction of protein functions. In order to compute the error, we use a binary crossentropy loss^33^ given as:

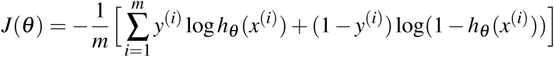

Finally, we train our classifier using the recent and highly successful ADAM classifier^34^ with the values for *β*_1_ as 0.9, *β*_2_ as 0.999 and the fuzz factor as *e*^−8^ while the learning rate was set to 0.001. We arrived at these values through random sampling from the hyperparameter space and performing cross validation on the results.

An important note that we would like to make about our model is regarding the issue of covariate shift in neural networks^35^. Internal covariate shift was a major issue in our initial experiments and we see this as the primary reason that deep learning has not yet been applied to the problem of protein function prediction. In typical machine learning problems, the problem of covariate shift arises when the distribution of real world data is different from that of training data^35^. In neural networks the learning is distributed across different layers. The output of each layer forms the input to the next layer. During backpropagation, the parameters of each layer are updated thus effectively changing the distribution of the inputs of the next layer. This is termed as the concept of internal covariate shift^36^. This can be tackled by normalizing the inputs of each batch^37^, which is given as 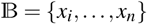. First we calculate the batch mean and variance and normalize each *x_i_*.

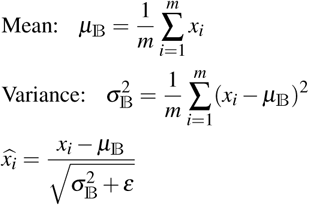

Here, *m* is the batch size which we set to 500 for this particular model. The layer has two trainable parameters *β* (shift parameter) and *γ* (scale parameter). The output values *y_i_* of a BatchNorm layer are computed as:

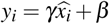

Without BatchNorm, our network failed to learn the patterns in the sequences and we were unable to achieve any reasonable accuracy even with very deep networks. With this normalization layer, we were able to improve the results even with shallower networks. In the next section, we benchmark the results we achieved by applying DeepSeq to the dataset of Homo sapiens species proteins.

## Results

A protein’s function generally describes molecular, cellular or phenotypic aspects that a protein is involved in, including how it interacts with other molecules (such as substrates, pathogens and other small compounds etc.). From the various proposed schemes to standardize the concept of protein function we select the most famous and widely used Gene Ontology (GO) classification scheme^38^ due to its desirable properties, e.g., wide coverage, disjoint categories, standardized hierarchical format etc. We operate on molecular function subconcept of the gene ontology, which is a hierarchical set of terms arranged in a Direct Acyclic Graph (DAG) structure depicting functional relationships among them. GO’s consistency across species and its widespread adoption make it suitable for large-scale computational studies.

The proposed scheme is benchmarked on proteins of a widely used modal organism of Homo sapiens species. We select five most frequent gene ontology target classes namely, ATP binding (GO:0005524), Metal ion binding (GO:0046872), DNA binding (GO:0003677), Zinc ion binding (GO:0008270) and Nucleic acid binding (GO:0003676) for training our model. The heterogeneous sub-ontology can be seen in Figure 4 with most frequent target classes highlighted through solid double border. The rationale for selecting these classes, besides the fact that they are the most frequently occurring ones, is that they come under the same molecular context i.e., binding, while preserving adequate heterogeneity.

**Figure 4.**
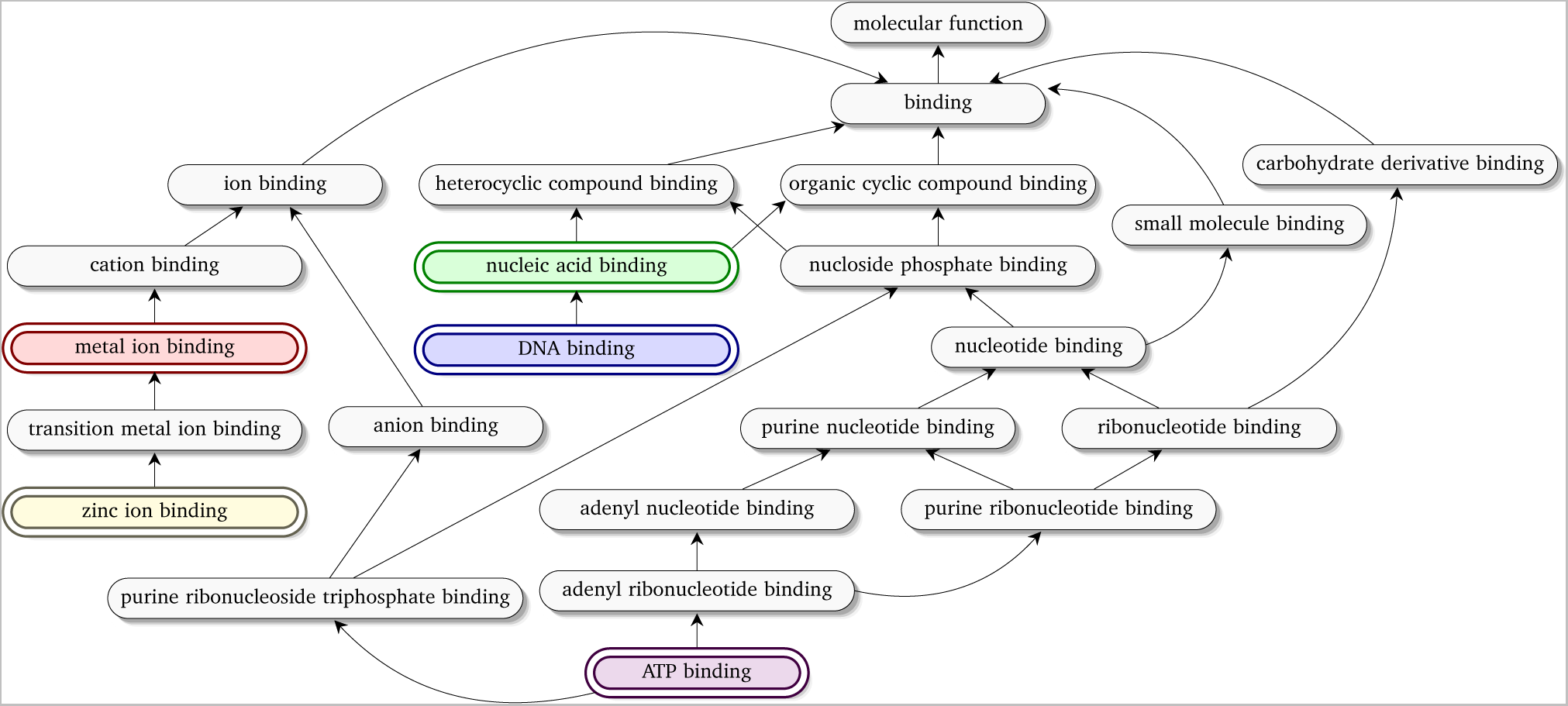
Five most frequently occurring GO terms (highlighted through double borders) belonging to molecular function branch of gene ontology.

The experiment was organized as follows: We retrieved a total of 72,945 proteins of Homo sapiens species from Uniprot database^39^. These proteins included both reviewed entries, called swiss-prot (sp), and unreviewed ones, called Tremble (tr). In order to maintain high prediction confidence we utilized only swiss prot (sp) enteries.

The length of a protein sequence i.e., no of residues, varies from protein to protein. In Homo sapiens species proteins, this length varies from 28 residues up to 34,358 residues. Only a small percentage (*≈* 2%) of sequences are above the length of 2,000 residues. Due to computational constraints and training overhead, in this work we consider sequences whose length is up to 2,000 residues. This limitation is only for training sequences. However, DeepSeq can make accurate predictions for larger sequences given one sub-sequence at a time.

We evaluate the predictive performance of our model in a 3-fold cross validation setting, i.e. the training consists of two third of randomly selected annotated proteins data while we test the model on remaining one third annotated proteins. For evaluation of our model, we calculate several performance measures, such as: precision, recall, accuracy, and F1 score defined as:

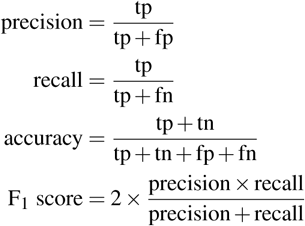

Where tp are the number of true positives, fp are the number of false positives, tn are the number of true negatives and fn are the number of false negatives. It is pertinent to mention here that by definition if a model predicts a parent gene ontology term of a true positive term, it is considered a true positive prediction but in reality, it is a less accurate prediction. Therefore, for our model, we compiled results by considering a term as true positive only if it exactly predicts that specific term.

A typical run of DeepSeq on a system with CUDA GPU K40, Core i7 CPU and 32GB RAM took approximately 75 minutes for 50 epochs of training. In Figure 5, we report the F1 score and relative loss for both test and training curves on 50 epochs of training. Our DeepSeq architecture achieves the best F1 score (with minimum loss) in the 43rd epoch. Further training tends to overfit the model i.e., it leads to a surge in the test loss and would perform poorly on newer, unseen data points. We therefore stop training at this point thus effectively employing the technique of early stopping^40, 41^.

**Figure 5.**
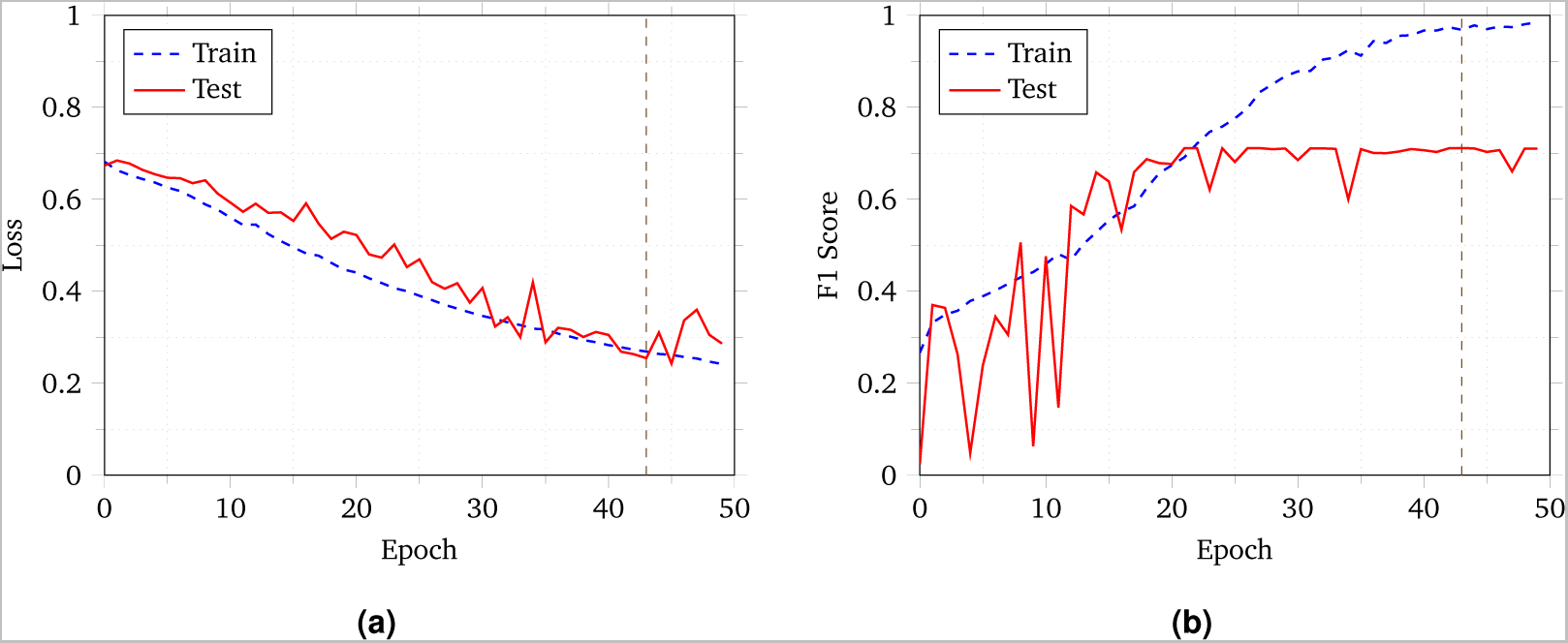
Learning curves in DeepSeq for Homo sapiens species proteins. Early stopping was employed at epoch 43 to prevent overfitting of data.

### Comparison with other methods

We comprehensively compare our model with the most widely used BLAST algorithm, which is based on homology based annotation transfer and utilizes only protein sequence information for function prediction. BLAST is a local alignment based search algorithm that finds hits among protein sequences (query and database sequences), based on local regions of similarity by utilizing a heuristic function. For Homo sapiens species proteins, we report accuracy, precision, recall and F1 score of BLAST algorithm as well as our method in Table 1. Clearly our method outperforms BLAST in all aspects. It is mainly because BLAST algorithm results in many false positives for proteins with multiple functions. We also observe that the downgraded performance of BLAST is attributed to two main factors: 1) due to heterogeneous nature of even very similar proteins and 2) BLAST’s complete annotation transfer when homology strikes with a high sequence similarity. Both these factors result in an increased number of false positives, because of which both precision and accuracy of BLAST drops significantly. On the other hand, DeepSeq is robust in annotation transfer as it considers not only the input sequence but also learns representations using training sequences. This enables it to reduce the false positive rate.

**Table 1.**
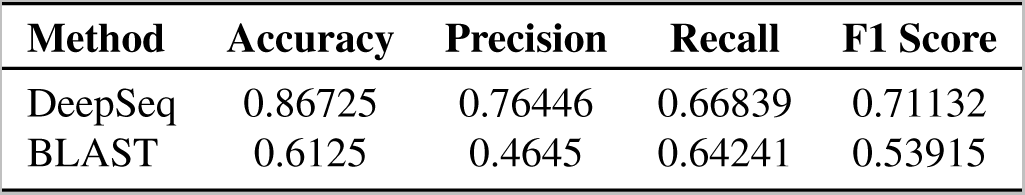
Comparison of accuracy, precision, recall, and F1 Score of our method with BLAST algorithm^8^.

### Effect of different parameters on DeepSeq architecture

In this section, we briefly elaborate the alternatives to our choices of parameters for DeepSeq, which resulted in sub-optimal results as well as those that we have not yet explored but form potential candidates for future studies. Initially, we set the filter size of each conv1d layer to 3 instead of 15. However, this small filter size was unable to find meaningful patterns in the sequences. Moreover, improvements to results became insignificant beyond filter size 15. Increasing the number of filters per layer from 250 and 100 also showed little improvements but led to difficulties in loading the model in memory. Similarly, increasing the number of layers was only possible by decreasing the number of filters per layer but did not lead to improvements in results.

Finally, we would also like to note that we have experimented with more complex model such as Long Short Term Memory (LSTM)^42, 43^ models and Gated Recurrent Units (GRUs)^44^. However, we were not able to achieve good results with these models primarily due to the long sequence sizes. Attention-based models^45^ might be useful in such a scenario but we are yet to explore these in detail. This forms an important and potentially fruitful future direction for this line of research.

### Localization and biological insight

In addition to the prediction of protein functions, another important aspect of our model is the ability to localize exact residue positions in the query sequence that are involved in a particular molecular activity, giving a deeper insight into understanding the function-residue relationships. In Figure 6, we show an example sequence for *DNA repair and recombination protein RAD54-like* protein (Uniprot ID: A0A087WTB0 HUMAN), that is annotated with ATP binding function.

**Figure 6.**
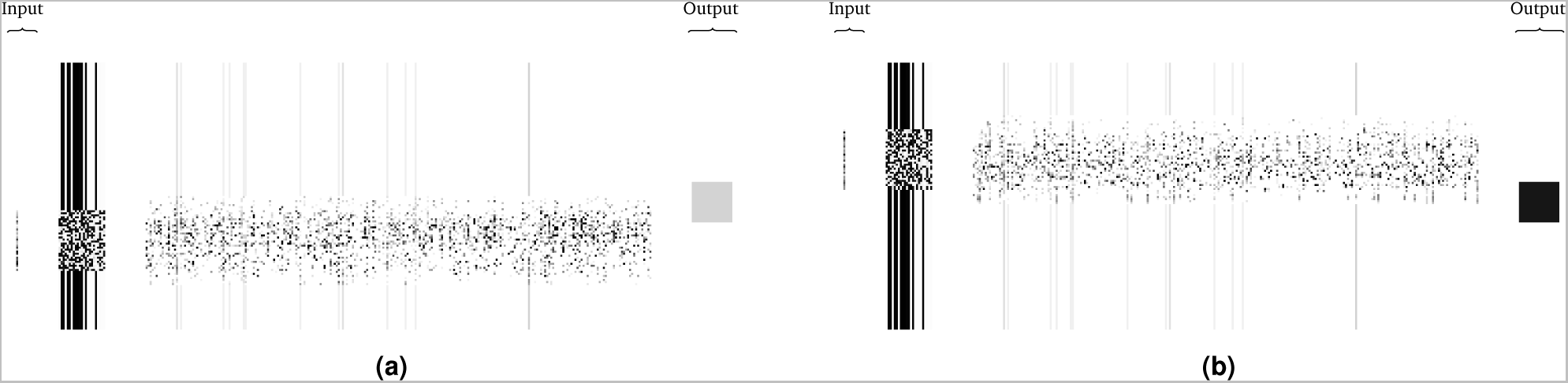
Protein localization through DeepSeq for the ATP binding function. We fix a window size (here, set to 20) and replace everything outside the window with 0s. This window is slid over the whole sequence and outputs of the network observed. If a positive prediction is made by DeepSeq, we associate the region of the window with the ATP binding function. Here, darker color signifies positive prediction.

The behavior of DeepSeq can be seen in Figure 6a when the input is a subsection of the sequence which is not responsible for ATP binding (GO ID: 0005524) function. The network’s output layer node responsible for ATP binding produces a low value, shown as a square in a lighter shade of gray. On the contrary, when the input is a different region of the input sequence (as shown in Figure 6b) that is responsible for ATP binding function, the output perceptron for ATP binding fires at a high value, shown as a dark square. Thus, DeepSeq has the ability to localize the exact sub-sequence of a protein that is responsible for a molecular activity. The localized sub-sequences may vary in size, and an important aspect of utilizing this variation could be to assess the reliability of existing function-motifs/function-domain associations in related biological databases.

## Discussion

Protein function is a concept that is elusive in nature and its meaning can vary in different biological contexts. The prediction of protein function is challenging due to a number of reasons. First, protein functions have impact at multiple levels e.g., a protein might be part of some biochemical activity at the molecular level and at the same time it affects metabolic pathways, cells, tissues, up to the entire organism. Second, proteins are heterogeneously multifunctional and highly chaotic in nature; in fact, more than 30% proteins in Swiss-Prot have multiple functions^46^. Third, the available function information is incomplete due to the fact that its experimental characterization is context dependent i.e., a single experiment is unlikely to cover all contextual conditions (such as temperature, interactor proteins, pH etc.) under which protein’s entire functional repertoire could be revealed. Lastly, in addition to being incomplete, existing function information in world wide databases are error prone because of experimental biases, interpretation or curation issues. Despite all these issues, automated algorithms are showing progress as evident from the most recent Critical Assessment of Function Annotation (CAFA-2) experiment.

Our proposed architecture – DeepSeq – is a fully automatic deep learning algorithm, which predicts functions of a protein using only sequence information that is available in abundance. The proposed architecture has been benchmarked, training it on five most frequent gene ontology target classes, namely ATP binding (GO:0005524), metal ion binding (GO:0046872), DNA binding (GO:0003677), zinc ion binding (GO:0008270) and nucleic acid binding (GO:0003676).

The extracted classes are heterogeneous (i.e., ontology leaf nodes are spread with many branches) and their accurate annotation is a challenge in itself for traditional classifiers because of two reasons: First, all the five terms come under the same context, which makes it difficult to find associated fingerprints in the input sequence to help in the characterization process. Second, the functional context becomes specific as we go deeper in the ontology and more information (in the form of features/representations) is needed to distinguish such target classes. However, DeepSeq was able to extract representations for five most frequent ontology classes and was remarkable in predicting each ontology class accurately.

### Prediction of functionally unknown proteins

We extracted 40,348 uncharaterized proteins (as of March, 2017), from Homo sapiens data set of the Uniprot database. Our method assigned plausible functions to 4,594 proteins, a vast majority (2,470) of which potentially belonged (verified through sequence homology) to one of five classes that we trained our algorithm on. The distribution of predictions of the five top functions can be seen in Figure 7 whereas the complete list of proteins and their predicted annotations is available (filename: `predictions-unknowns.txt`) in the Github repository at http://github.com/recluze/deepseq. An important thing to note here is that the algorithm is not constrained to the size of the input sequence and annotated uncharacterized proteins include sequences whose length is larger than the trained sequences (i.e., *>*2000 residues).

**Figure 7.**
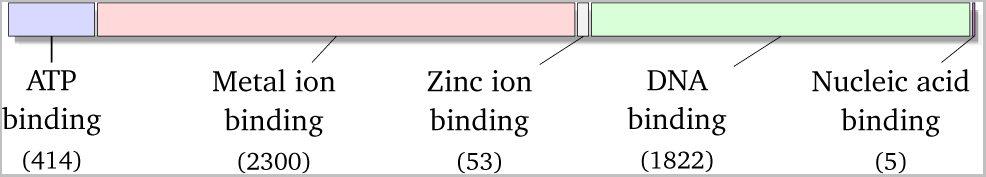
Distribution of functions predicted for unknown proteins.

### An example annotation case study

In this section we illustrate a working example of a known protein from the Homo sapiens species, highlighting DeepSeq’s ability to accurately predict heterogeneous GO categories. We chose the *2’-5’-Oligoadenylate Synthase 1* (short name: *2-5A Synthase 1*) protein for our analysis, which is a 400 amino acid long protein, with 3 experimentally verified InterPro domains^47^. The *2-5A synthase 1* protein has a fully reviewed annotation in Uniprot’s swiss-prot database, with experimentally verified functions, which make it a suitable candidate to demonstrate the working of our model. The *2-5A synthase 1* protein is positively annotated with the gene ontology terms for which DeepSeq is trained i.e., ATP binding (GO:0005524), metal ion binding (GO:0046872), and zinc ion binding (GO:0008270). The involvement of this protein in multiple gene ontology categories makes prediction of its functions a challenging task. However, as described below, DeepSeq is able to automatically extract features and infer accurate functions as well as perform localization using only the protein sequence.

When the *2-5A synthase 1* protein’s sequence is given as input to our model, the 3 output layer neurons out of five responsible for ATP binding, metal ion binding, and zinc ion binding functions fire a positive value showing true positive predictions for all three trained functions. However, the conceptual novelty is not just the prediction of functions. Another important aspect of our deep learning architecture is the exact localization of features/fingerprints in the input sequence that are responsible for a particular activity. We verified as well as compared each molecular function’s location discovered by our model with associated InterPro domains.

Our example protein has three InterPro domains namely: 2-5-oligoadenylate synthetase, N-terminal (IPR006116), Polymerase, nucleotidyl transferase domain (IPR002934) and 2’-5’-oligoadenylate synthetase 1, domain 2/C-terminal (IPR018952). The IPR006116 domain starts at residue 17 and ends at residue no. 105, IPR002934 domain starts at residue 37 and ends at residue no. 123 and IPR018952 domain starts at residue 164 and ends at residue no. 344. Each domain is associated with a number of molecular function activities. For example IPR006116 domain is associated with RNA binding (GO:0003723), ATP binding (GO:0005524), transferase activity (GO:0016740) and immune response (GO:0006955); thus a protein known to have this domain potentially participates in any of these activities.

In general, because of multiple molecular activity associations, the presence of a domain in a protein sequence doesn’t exactly localizes residues for a particular molecular activity. DeepSeq, on the other hand, is able to locate more specific residue positions. (A schematic illustration is added in Figure 8.) The network predicted ATP binding (from residue position 21 to 51), metal ion binding (from residue position 38 to 58), and zinc ion binding (from residue position 56 to 86), when given 20 residues as input at a time for the first two and 30 residues at a time for the last one.

**Figure 8.**
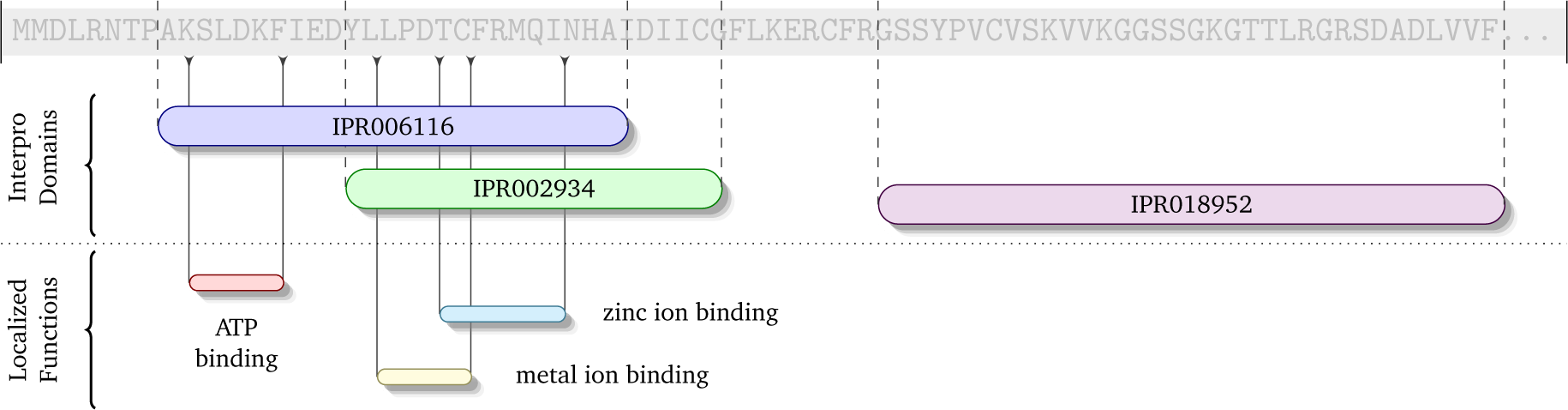
Schematic view of the annotation case study for protein 2’-5’-Oligoadenylate Synthase 1. Not to scale.

### Transfer learning

Finally, we describe another important byproduct of DeepSeq that has significant scope beyond the work presented in this paper. One of the important aspects of applying deep learning to a problem is the transferability of extracted features to other problems. This is termed as ‘transfer learning’ and is a highly significant area of ongoing research^48, 49^. Once we have successfully applied our architecture (depicted in Figure 3) to the problem of function prediction, the network is able to not only predict the functions successfully but in the process, it has also learned to extract representative features. These features of the input sequence are used by the classifier on the last layer of the network.

The concept of transfer learning dictates that we can remove the last layer from the network, freeze the weights of the initial layers and affix a different layer to the end of the network. If this new layer is able to perform another type of classification, the features learned in the earlier phases can be used to solve a different type of problem with only minimal training (i.e. learning the weights of just the last layer). Since we provide the full network architecture of our model as well as the learned weights as open source, we envision that our model, coupled with transfer learning, can be used to solve a host of other problems such as prediction of 2D and 3D structure of a protein.

Taking a step even further, we may combine two or more instances of our feature extraction layers to two or more distinct protein sequences. We can then merge the outputs of these feature extraction layers and add a classifier to the merged output. Through this simple trick, DeepSeq can make predictions about potential bioinformatics problems involving *multiple* protein sequences, such as the prediction of protein-protein interactions. All of this forms part of our ongoing research work and we hope the community will be able to benefit from our model on all these fronts.

## Conclusion

In this paper, we have presented DeepSeq – a novel deep learning architecture that takes the sequence of a protein as input and accurately predicts its functions without relying on any supporting information. An important aspect in the prediction process is the ability to automatically extract representations from the input protein sequence without the need for a human expert. We achieved significant improvements in precision, accuracy and F1 score for the proteins of Homo sapiens species. Besides the prediction of protein functions, we have also used the trained network to localize region of the sequence that is involved in a particular molecular activity. An example with query sequence and localized functions is also presented. In addition, the trained network parameters can also be used for transfer learning (i.e., learning a *different* concept), say, predicting the potential 3D conformations of a protein given its sequence.

Thus, we have shown that DeepSeq can extract meaningful information from the input sequence that can then be used to solve a myriad of complex problems without human intervention. We envision that future efforts in applying DeepSeq and its variants will lead to ground breaking successes in the field of bioinformatics just as deep learning has revolutionized other fields such as computer vision, natural language processing and machine translation.

## Acknowledgements

Hafeez Ur Rehman’s contribution in this work was partially supported by Grant Number: 21-915/SRGP/R&D/HEC/2016 by the HEC. The execution of our deep learning experiments was made possible by the gracious contribution of a Tesla K40 GPU by NVIDIA Corporation. The contents of this paper are not necessarily endorsed by the funding agencies.

## Author contributions

M.N. conceived the idea of using deep learning for Bioinformatics, H.R. provided domain knowledge and structured the problem. Both M.N. and H.R. wrote the code for the experiments. G.P. contributed with running experiments and analyzed results. A.B. helped analyze results and formalize the details of discussion. All authors contributed in manuscript preparation and review.

## Footnotes

1 Instructions for executing the code are in the included README file.

